# Developing Gene-Specific Meta-Predictor of Variant Pathogenicity

**DOI:** 10.1101/115956

**Authors:** Anna Rychkova, MyMy C. Buu, Curt Scharfe, Martina I. Lefterova, Justin I. Odegaard, Iris Schrijver, Carlos Milla, Carlos D. Bustamante

## Abstract

Rapid, accurate, and inexpensive genome sequencing promises to transform medical care. However, a critical hurdle to enabling personalized genomic medicine is predicting the functional impact of novel genomic variation. Various methods of missense variants pathogenicity prediction have been developed by now. Here we present a new strategy for developing a pathogenicity predictor of improved accuracy by applying and training a supervised machine learning model in a gene-specific manner. Our meta-predictor combines outputs of various existing predictors, supplements them with an extended set of stability and structural features of the protein, as well as its physicochemical properties, and adds information about allele frequency from various datasets. We used such a supervised gene-specific meta-predictor approach to train the model on the *CFTR* gene, and to predict pathogenicity of about 1,000 variants of unknown significance that we collected from various publicly available and internal resources. Our *CFTR*-specific meta-predictor based on the Random Forest model performs better than other machine learning algorithms that we tested, and also outperforms other available tools, such as CADD, MutPred, SIFT, and PolyPhen-2. Our predicted pathogenicity probability correlates well with clinical measures of Cystic Fibrosis patients and experimental functional measures of mutated CFTR proteins. Training the model on one gene, in contrast to taking a genome wide approach, allows taking into account structural features specific for a particular protein, thus increasing the overall accuracy of the predictor. Collecting data from several separate resources, on the other hand, allows to accumulate allele frequency information, estimated as the most important feature by our approach, for a larger set of variants. Finally, our predictor will be hosted on the ClinGen Consortium database to make it available to CF researchers and to serve as a feasibility pilot study for other Mendelian diseases.

## Introduction

The advent of next-generation sequencing that is quick, accurate, and affordable has promised to usher in a new era of genomic medicine. However, a critical issue facing the development of sequencing-based tests is the interpretation of novel genetic variants in terms of their probability of causing disease. This is a particularly pressing problem with so-called “clinically relevant genes”, including the cystic fibrosis transmembrane conductance regulator (*CFTR)* gene, for which DNA changes are known to impact phenotype, but for which the map of how each genotype affects the clinical phenotype is incomplete. Differentiating “benign” from “pathogenic” genetic variants is challenging, and often physicians are left with the unsatisfying and inconclusive result that their patient carries a “Variant of Unknown Significance” (VUS).

Despite recent advances in applying machine learning techniques to problems in biomedicine, existing computational approaches to variant classification all suffer from low overall accuracy rates. For example, SIFT^1^ and PolyPhen-2^2^ are among the most widely used algorithms, but each has an accuracy of less than 70%^3^. Their poor performance limits the clinical utility of these tools in determining whether a novel genetic variant is actually related to the disease of interest. We aimed to improve the performance of computational interpretation tools by developing a gene-specific metapredictor, focusing on the *CFTR* gene, which combines information from the most promising available tools supplemented with protein structure and stability features, physicochemical properties of mutated residues, and allele frequency information.

To develop this computational model, we focused our analyses on variants in the coding region of the *CFTR* gene. Despite recent progress in both sequencing and analysis techniques, interpreting the functional effect of variants in non-coding regions remains problematic due to insufficient training data. Therefore, to maximize the prediction capability of our model, we are initially focusing solely on the variants that are most likely to be relevant in terms of disease association, due to their relatively clearer relationship to protein structure.

Many bioinformatics methods have been developed for predicting the effect of missense mutations, which vary by the number of features included and the type of machine learning algorithm employed. The most advanced tools typically rely on amino acid sequence, protein structure, and evolutionary conservation for their prediction. For example, while SIFT relies solely on conservation, measured via multiple sequence alignment, PolyPhen-2 includes both sequence and structure-based features for prediction. The structure-based features in this context are used to describe the physical environment of the mutation, and include predictors such as solvent-accessible surface area, hydrophobic propensity, and the “mobility” of the atom. Another missense pathogenicity predictor, MutPred^4^, uses a much larger set of structural parameters, including secondary structure, stability and intrinsic disorder, transmembrane and coiled-coil structure. In addition, MutPred utilizes functional properties of the protein, such as sites of post-translational modification, catalytic and DNA-binding residues. MutPred outperforms SIFT by 7% in the area under the ROC curve (AUC), and, more importantly, in addition to pathogenicity score can provide information about the molecular basis of the disease.

While SIFT, PolyPhen-2 and MutPred are trained using data from across the genome, gene-specific pathogenicity prediction methods have also been developed. For example, Masica et al.^5^ created a *CFTR*-specific prediction algorithm called Phenotype-Optimized Sequence Ensemble (POSE). In contrast to methods utilizing multiple sequence alignment, POSE tries to iteratively construct an optimized sequence ensemble based on the performance of the scoring function, which uniquely integrates evolutionary conservation with physicochemical properties of the amino acids (such as charge, presence of aromatic or aliphatic group, hydrogen bond donor or acceptor, and signals for glycine and proline residues). POSE achieves a performance of 84%, as measured by AUC on a training set of 103 *CFTR* variants, and importantly, the method displayed improved specificity when compared to tools trained genome-wide, implying a higher accuracy potential for methods trained on single genes.

Combining existing methods into a single predictor has proven to yield increased accuracy^6,7^. Successful examples of such meta-predictors, therefore, suggest that the separate methods used for prediction of variant-disease associations are orthogonal, and represent different biologically relevant relationships. The advantage of the machine learning classifier is its ability to integrate these orthogonal measures to identify predictive signatures of pathogenicity. Thus we are employing such a combination strategy in developing our *CFTR*-specific meta-predictor.

In addition to combining outputs from several existing prediction tools, we are also adding other useful features into out meta-predictor. Importantly, we integrate protein stability measures into our pathogenicity predictor. Protein stability is a fundamental property that affects function, activity, and regulation of biomolecules. Conformational changes are required for many proteins’ function, implying that conformational flexibility and rigidity must be finely balanced. Incorrect folding and decreased stability are two of the major consequences of missense mutations, which can lead to disease. Protein stability is measured by the folding free energy change upon mutation, which is calculated as the difference in free energy between the folded and unfolded protein states^8^. Therefore, the estimated folding free energy change for each variant should give valuable information about the functional consequence of missense mutations.

By restricting our method to one particular gene, we are trying to take advantage of the protein structure information and extract from it features uniquely relevant for the CFTR protein. Unfortunately, the full protein structure for CFTR has not yet been solved with X-ray crystallography. However, a few homology models have been built based on the available crystal structure of the nucleotide-binding domain and the homologous ABC transporter, Sav1866^9,10^. Structural parameters that have been tightly fitted to the CFTR protein, as well as inferred changes in physicochemical properties induced by amino acid substitution, are valuable features that help to increase the overall method performance.

## Materials and Methods

### Variants data collection

We utilized various data sources, both publically available and internal, to collect known protein coding variants in the *CFTR* gene (**Table 1**). Public data sources include The Clinical and Functional TRanslation of *CFTR* (CFTR2)^11^, the database of Genotypes and Phenotypes (dbGaP)^12^, and the Exome Aggregation Consortium (ExAC)^13^. Internal resources include datasets obtained from the Stanford Cystic Fibrosis Center^14^, and the Stanford Molecular Pathology Laboratory^15,16^. In addition we also included variants used for training and testing the previously described POSE method, which was trained directly on *CFTR^5^.* Overall, 1,899 protein coding *CFTR* variants have been collected, of which the majority (>60%) are missense variants (**Fig. 1A**). Clinical significance (pathogenic, benign, variant of unknown significance (VUS)) was merged from different sources, and conflicting entries (reported as pathogenic by one source and benign by other) were considered as VUSs. Since the ExAC database does not report variants’ pathogenicity, all the *CFTR* variants from ExAC were considered as VUSs. As expected, only a small portion of collected variants had pathogenicity evidence (14% pathogenic, 7% benign) (**Fig. 1B**), with the majority (~80%) having unknown significance.

**Table 1.** *CFTR* variants data collection.

| **datasource** | **# of protein-coding variants** | **# of unique variants for this datasource** |
| --- | --- | --- |
| The Stanford Molecular Pathology Laboratory | 1267 | 726 |
| The Exome Aggregation Consortium (ExAC) | 937 | 559 |
| POSE (*CFTR*-specific pathogenicity prediction algorithm) | 240 | 37 |
| The Clinical and Functional Translation for *CFTR* (CFTR2) | 171 | 4 |
| The Stanford Cystic Fibrosis center | 80 | 5 |
| The database of Genotypes and Phenotypes (dbGaP) | 53 | 9 |
| **Total:** | **1899** |  |

Variants data have been collected from six data sources, including external publicly available data and internal data from Stanford-affiliated laboratories. Total number of unique variants collected is 1,899.
\*The Stanford MPL database that we used include variants identified during clinical CF testing at the lab, and laboratory-curated *CFTR* variants from various public datasources, including the CF Mutation Database, the CFTR2 database, National Center for Biotechnology Information's dbSNP, and Ensembl.

**Figure 1.**
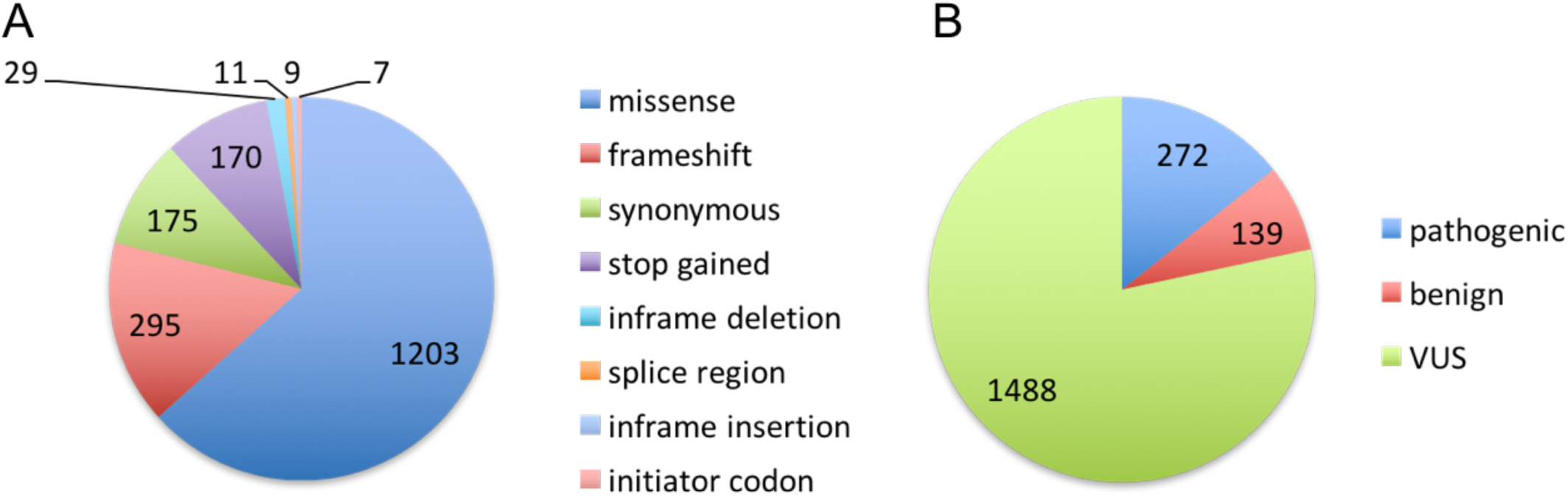
Distribution of 1,899 collected protein coding CFTR variants by mutation type and clinical significance. (A) Missense variants represent the largest class of protein-coding variants in *CFTR.* (B) Majority of variants are classified as VUSs.

The full dataset of 1,899 *CFTR* variants with clinical significance from corresponding sources can be found on GitHub (https://github.com/rychkova/CFTR-MetaPred).

### Variants annotation

Our meta-predictor is built by combining outputs from a number of the available prediction tools and supplementing them with information extracted from protein structure and allele frequency (**Table 2**). From the variety of pathogenicity prediction tools available we considered those based on evolutionary conservation only (PROVEAN^17^, SIFT^1^, PANTHER^18^), and those based on some additional structural information as well (PolyPhen-2^2^, MutPred^4^, CADD^19^, POSE^5^). Information regarding individual allele counts and overall sample sizes of the different studies was combined and converted to allele frequencies of variants. We used *density* function in R with the default Gaussian smoothing kernel to estimate the probability density function from the allele frequencies. Given the importance of protein stability for proper cellular function, we also incorporated predictors of folding free energy change into our model (Eris^20^, PoPMuSiC^21^, FoldX^22^). We used two available homology models of CFTR protein^9,10^ for each of the stability predictor, which gave rise to six total stability features.

**Table 2.**
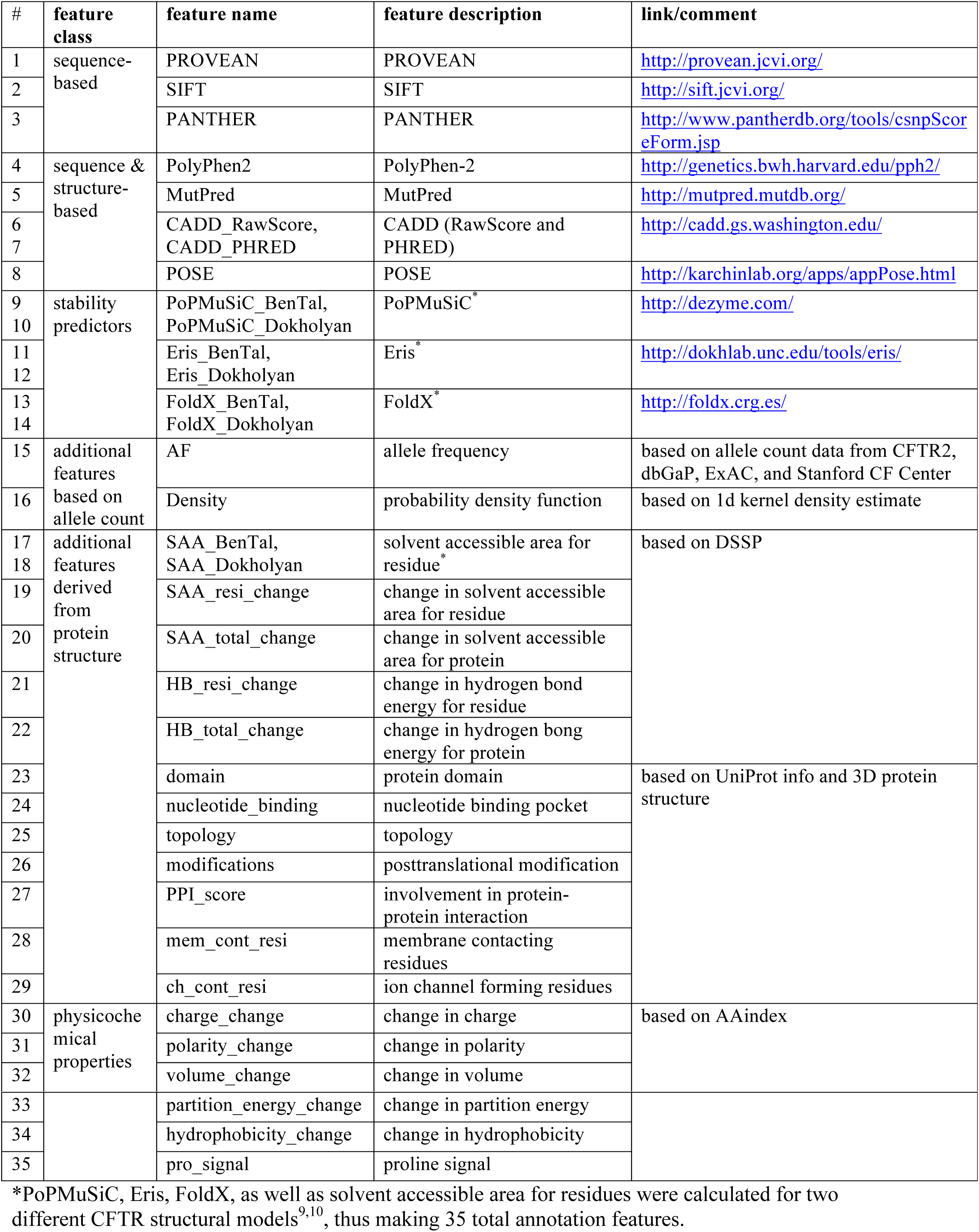
Features used for training machine learning algorithms.

We further created several structural parameters based on the information in UniProt^23^ and 3D CFTR protein structure, such as protein domains (extracellular loops, nucleotide binding domain 1 or 2 (NBD1 or NBD2), transmembrane domain 1 or 2 (TMD1 or TMD2), R domain), nucleotide binding residues, topology (cytoplasmic, transmembrane, or extracellular protein parts), regions of posttranslational modification (phosphorylation, glycosylation, palmitoylation, or ubiquitination sites), and involvement in protein-protein interaction (PPI_score). Our PPI_score for each residue is based on the number of times each residue is present in the motifs known to be important for protein-protein interaction and CFTR regulation. Information about protein-protein interaction motifs known for CFTR is based on the literature, and summarized in **Table S1**. On top of these we added information about membrane contacting residues by building a simplified membrane model around the protein (using Coarse Grained model building tool in Molaris^24^), and selecting neighboring to membrane atoms residues in PyMol^25^. Similarly, we created a feature with channel contacting residues, by inserting a straight helix into the channel and selecting neighboring residues in PyMol.

We used DSSP tool^26^ to calculate change in solvent accessible area and hydrogen bond energy of the full protein as well as single residue upon mutation. Structural models of all the 1,210 mutant proteins (missense variants plus initiator codon variants) were obtained with Eris program^20^ using the default fixed-backbone method. Change in several physicochemical properties of residue due to mutation was estimated based on the information in the AAindex dataset^27^ (charge, polarity, volume, partition energy, hydrophobicity, proline signal (mutation to/from proline)).

The full dataset with 35 annotation features collected for all the *CFTR* missense variants can be found on GitHub (https://github.com/rychkova/CFTR-MetaPred).

### Machine learning model training

To build the machine learning model, we utilized the statistical software program R with the library package caret^28^. To find the best performing algorithm, we tested several available methods: regularized logistic regression (GLM), regularized discriminant analysis (RDA), support vector machine (SVM), stochastic gradient boosting (tree boosting method) (GBM), and random forest (RF). The description of all the methods can be found in ref^29^. Of the 1,210 missense and initiator codon variants we annotated with 35 features, 295 unique variants had known clinical significance (161 pathogenic, 134 benign). We performed data preprocessing step by converting all the categorical features into numeric values, converting all the values into Z-scores, and imputing data with KNN method. It should be noted that a considerable amount of missing allele frequency data did not allow for a KNN imputation of this category, thus we used a dummy value of -1 for all the missing allele frequencies. We divided our dataset into training and testing sets with a ratio of 70/30. Five different models were built on the training set using five-fold cross validation for resampling, and the performance was measured on the test set. We have also estimated the performance of all the separate 35 features on the training set and compared it with the machine learning models.

## Results

Performance of all the five machine learning models we built and the 35 separate features can be found in **Tables 3** and **4**, respectively. Of the five models tested, RF showed the highest accuracy (77%). Based on the AUC values, RF model outperformed all the other machine learning models (AUC_RF_ = 85%) (**Fig. 2A**), and it also improved over other popular tools, such as CADD (AUC_CADD_._Rawscore_ = 70%), SIFT (AUC_SIFT_ = 63%), and PolyPhen-2 (AUC_PolyPhen2_ = 60%) (see **Fig. 2B** and **Table 4**). Out of the 35 separate features, AF showed the best performance (AUC_AF_ = 73%) (**Fig. 2B**). AF, Density, MutPred, POSE, and SIFT were selected as the most important features by the RF model (see **Fig. 3** and **Table S2**). Interestingly, when looking at the features by their class (as defined in **Table 2**), features based on allele count (AF and Density) seem to be the most significant ones (**Table S2**), followed by sequence & structure-based predictors (MutPred and POSE). SIFT was selected as the most significant one out of three sequence-based predictors we used (SIFT, PROVEAN, PANTHER). With regards to features derived from protein structure, protein topology (transmembrane helix, cytoplasmic or extracellular domain) and information about number of protein-protein interactions the residue participates in (PPI_score) showed higher importance, than other features calculated by DSSP for mutated protein models (HB_resi_change, HB_total_change, SAA_resi_change, SAA_total_change). From six physicochemical property features we derived from AAindex database (volume_change, polarity_change, partition_energy_change, hydrophobicity_change, charge_change, pro_signal), change in residue volume upon mutation seems to be the most important one.

**Table 3.** Performance measures for the five machine learning algorithms tested.

| method | accuracy | sensitivity<br>(TPR=TP/P) | specificity<br>(TNR=TN/N) | AUC |
| --- | --- | --- | --- | --- |
| RF (random forest) | 0.77 | 0.75 | 0.80 | 0.85 |
| GBM (gradient boosting method) | 0.73 | 0.65 | 0.83 | 0.83 |
| GLM (generalized linear model) | 0.55 | 1.00 | 0.00 | 0.82 |
| SVM (support vector machine) | 0.68 | 0.71 | 0.65 | 0.79 |
| RDA (regularized discriminant analysis) | 0.68 | 0.71 | 0.65 | 0.77 |

TPR – true positive rate, TP – number of true positives, P – total number of positives (true positives + false positives), TNR – true negative rate, TN – number of true negatives, N – total number of negatives (true negatives + false negatives). Accuracy = (TP+TN)/(P+N).

**Table 4.** Performance measures for separate predictors.

| feature | accuracy | sensitivity<br>(TPR=TP/P) | specificity<br>(TNR=TN/N) | AUC | RF AUC<br>improvement<br>, % |
| --- | --- | --- | --- | --- | --- |
| AF | 0.73 | 0.69 | 0.78 | 0.73 | 11.9 |
| CADD_PHRED | 0.64 | 0.75 | 0.50 | 0.71 | 14.1 |
| CADD_RawScore | 0.63 | 0.60 | 0.65 | 0.70 | 15.0 |
| MutPred | 0.60 | 0.83 | 0.33 | 0.69 | 15.5 |
| PROVEAN | 0.65 | 0.69 | 0.60 | 0.66 | 19.0 |
| PoPMuSiC_BenTal | 0.63 | 0.71 | 0.53 | 0.63 | 21.9 |
| SIFT | 0.57 | 0.92 | 0.15 | 0.63 | 22.1 |
| SAA_resi_change | 0.56 | 0.88 | 0.18 | 0.62 | 22.9 |
| PANTHER | 0.60 | 0.73 | 0.45 | 0.62 | 23.0 |
| PolyPhen2 | 0.57 | 0.83 | 0.25 | 0.60 | 25.1 |
| SAA_Dokholyan | 0.61 | 0.69 | 0.53 | 0.59 | 26.2 |
| FoldX_BenTal | 0.53 | 0.92 | 0.08 | 0.59 | 26.2 |
| POSE | 0.52 | 0.67 | 0.35 | 0.58 | 26.8 |
| charge_change | 0.53 | 0.85 | 0.15 | 0.58 | 27.3 |
| SAA_total_change | 0.53 | 0.85 | 0.15 | 0.57 | 27.5 |
| volume_change | 0.50 | 0.75 | 0.20 | 0.57 | 27.6 |
| ch_contact_resi | 0.55 | 1.00 | 0.00 | 0.57 | 27.6 |
| domain | 0.55 | 0.85 | 0.18 | 0.57 | 28.1 |
| topology | 0.57 | 0.75 | 0.35 | 0.56 | 29.4 |
| PPI_score | 0.55 | 1.00 | 0.00 | 0.55 | 29.9 |
| FoldX_Dokholyan | 0.55 | 0.96 | 0.05 | 0.55 | 30.1 |
| hydrophobicity_change | 0.53 | 0.83 | 0.18 | 0.55 | 30.1 |
| PoPMuSiC_Dokholyan | 0.47 | 0.79 | 0.08 | 0.54 | 30.4 |
| mem_contact_resi | 0.58 | 0.98 | 0.10 | 0.54 | 30.9 |
| partition_energy_change | 0.50 | 0.90 | 0.03 | 0.54 | 31.3 |
| pro_signal | 0.55 | 1.00 | 0.00 | 0.53 | 32.2 |
| Eris_Dokholyan | 0.55 | 1.00 | 0.00 | 0.52 | 32.7 |
| Density | 0.55 | 1.00 | 0.00 | 0.52 | 33.1 |
| nucleotide_binding | 0.55 | 1.00 | 0.00 | 0.51 | 34.1 |
| modifications | 0.55 | 1.00 | 0.00 | 0.51 | 34.3 |
| polarity_change | 0.55 | 1.00 | 0.00 | 0.49 | 35.5 |
| HB_resi_change | 0.55 | 0.98 | 0.03 | 0.48 | 36.6 |
| HB_total_change | 0.55 | 0.98 | 0.03 | 0.48 | 36.6 |
| Eris_BenTal | 0.52 | 0.92 | 0.05 | 0.48 | 36.6 |
| SAA_BenTal | 0.55 | 1.00 | 0.00 | 0.47 | 37.7 |

**Figure 2.**
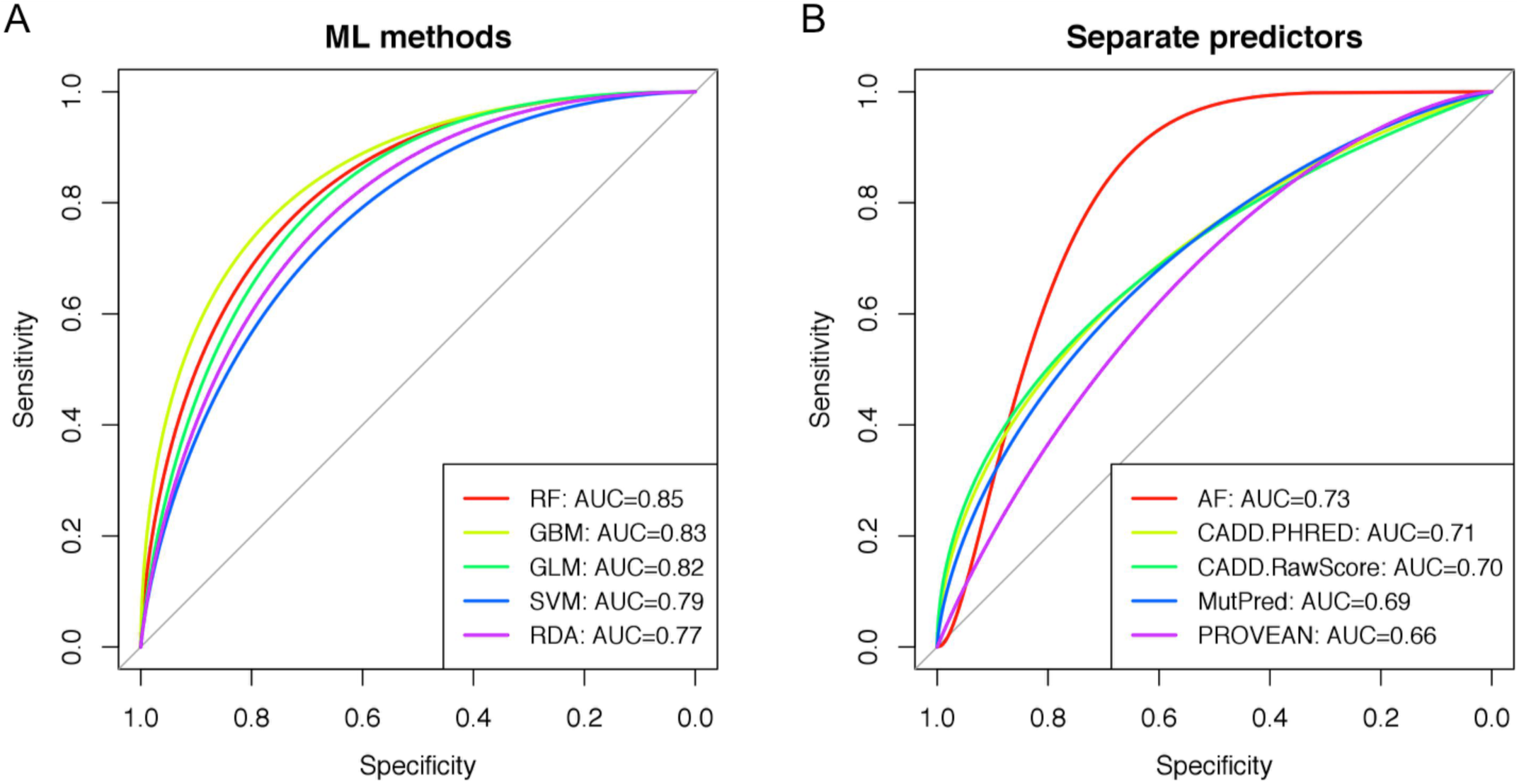
Performance comparison for ML algorithms tested and separate predictors used. (A) ROC curves for the five ML algorithms tested and their corresponding AUC values. Tree-based methods (GBM and RF) showed the highest performance, with the best AUC_RF_=0.85. (B) ROC curves for the separate predictors used for training. Only five best predictors out of 35 shown for clarity. AF predictor showed the highest AUC_AF_=0.73 in compare to other tools. AUC values for all 35 features are in **Table 4**

**Figure 3.**
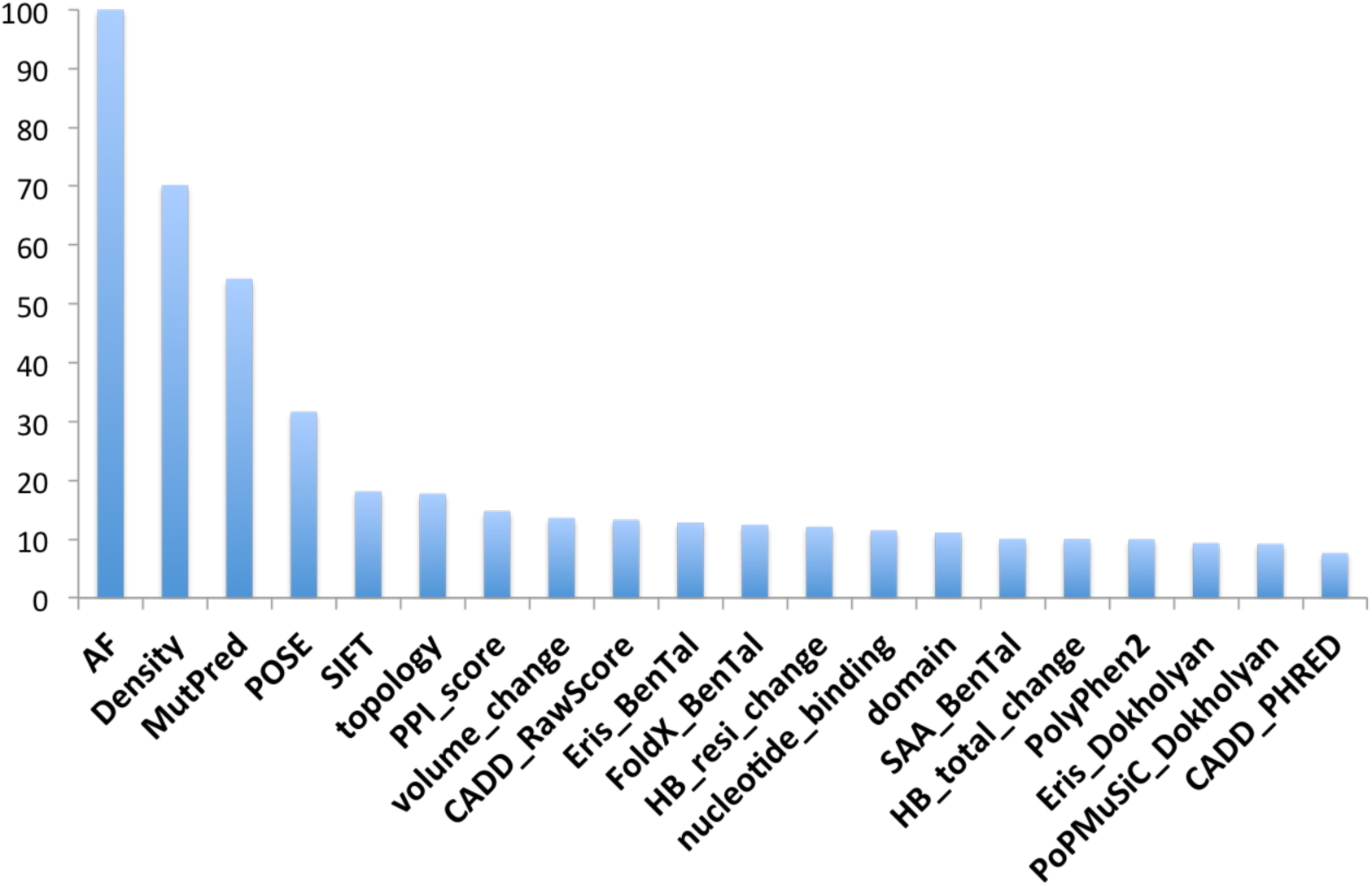
Features importance based on the RF model. Only top 20 features are shown. Values are scored from 0 to 100. Data for all 35 features are in **Table S2.**

To confirm our predictor’s validity, we have also examined how predicted pathogenicity probability correlates with existing clinical and functional data. We used previously measured mean chloride conductance values^30^ (**Fig. 4A** and **Table S3**) and sweat chloride data collected on patients at The Stanford CF Center (**Fig. 4B** and **Table S4**). The sweat chloride correlation analysis was restricted to patients heterozygous for p.F508del to reduce the variability due to different allele combinations. Both characteristics correlate well with the pathogenicity scores obtained using our RF classifier, with chloride conductance, which is a more direct measure of channel function, displaying the higher correlation coefficient (R^2^=0.44). Mean chloride conductance values, as well as mean sweat chloride concentration values are listed in **Tables S3** and **S4**, respectively.

**Figure 4.**
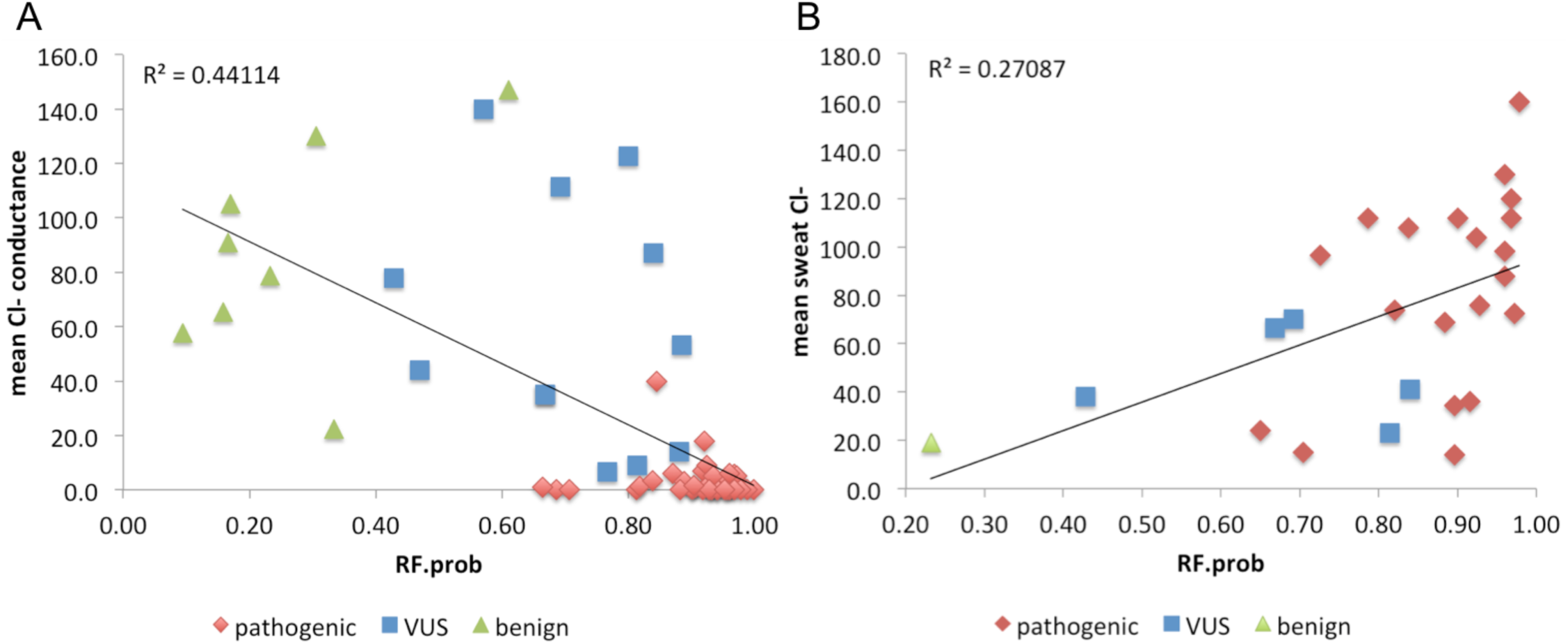
Correlation of pathogenicity prediction with experimental data. (A) Mean Cl^-^conductance values were taken from Sosnay et al.^30^. (B) Mean sweat Cl^-^ conductance based on the data from The Stanford CF Center. Values for patients heterozygous for F508del mutation only were used. Pathogenicity probability is based on the predictions from RF model.

Random forest pathogenicity probabilities along with predicted clinical significance categories for all the 1,210 missense *CFTR* variants can be found at GitHub (https://github.com/rychkova/CFTR-MetaPred
).

## Discussion

It has recently been recognized widely^31–34^ that computational predictors alone will not be able to reach satisfactory accuracy for direct use in the clinic, and both *in vitro* and *in vivo* functional studies are important to supplement the *in silico* predictions. Recognizing the importance of continuous-valued experimental quantitative measurements, rather than binary traits, Masica et al.^31^ extended their previously developed POSE method by including endophenotypic data from six clinical and functional assays. Their ePOSE (endoPhenotype-Optimized Sequence Ensemble) approach allows prediction of quantitative phenotypes associated with cystic fibrosis disease severity for missense variants in CFTR NBDs. Another study by Starita et al.^32^ explored the use of massively parallel experimental assays to measure the effect of nearly 2,000 missense substitutions in the RING domain of BRCA1 on its E3 ubiquitin ligase activity and its binding to the BARD1 RING domain. Model generated on the resulted scores was able to predict the capacities of full-length *BRCA1* variants, and outperformed widely used biological-effect prediction algorithms.

It should be noted that meta-predictor described here could easily be supplemented with functional data collected in high-throughput. *In vitro* functional measurements or even *in vivo* clinical phenotypes could be added as extra features during the model building step. With respect to CFTR protein, functional features like ion conductance, protein translation to the cell surface, and mRNA stability might be measured experimentally and added to the predictor for its overall performance improvement. Clinical phenotypes (like sweat chloride concentration, pancreatic sufficiency status, growth parameters, rate of first acquisition of *Pseudomonas aeruginosa* in the first year of life, and persistent colonization with *P. aeruginosa*) that are used in the clinic to help inform disease liability and penetrance of uncategorized mutations^35,36^, can be utilized as additional features in the prediction model as well. While our newly developed meta-predictor is not intended to uncover the molecular mechanisms of pathogenicity, it can help in prioritizing novel and rare genomic variations, identified through sequencing, for future functional studies. In addition, additional functional data may help to suggest potential cellular mechanism of the disease and allow for more accurate selection of specific therapies, as well as identify patients suitable for particular clinical trials.

The predictor described here could only be used to estimate a pathogenicity probability of missense variants, and its extension to other types of variants is somewhat less straightforward. The issue arises from the limited applicability of the predictors and structural features that we have utilized to build the model to classes of variants outside the missense category. In particular, only the methods PROVEAN and CADD can be applied to insertions and deletions, while nonsense, splicing, and synonymous mutations can be assessed by CADD only. To extract structural features, the structure of the mutated protein must be available, which is problematic for insertions and deletions that can cause large changes in structure. Moreover, the small set of TPs and TNs available for non-missense variant types limits our ability to train a similar meta-predictor, though allele frequency could in principle be used for any variant type. This again highlights the importance of additional functional measurements, which can be used alone or in combination with a few available computational features to establish the pathogenic status of all other types of variants.

By combining multiple levels of knowledge about *CFTR* structure and function, and training the machine learning model on the set of known pathogenic and benign variants, we created a *CFTR-*specific pathogenicity predictor tool of higher accuracy, which we hope may aid in interpreting and prioritizing *CFTR* variants, and be further evaluated by functional studies. This model’s predictions will be hosted on the ClinGen Consortium database, to make it easily available to other CF researchers and to demonstrate the feasibility of such an approach for a variety of Mendelian diseases. Overall, this report can be used as a description and model of the general strategy for developing a pathogenicity predictor of improved accuracy, so that feasibility of similar approaches may be evaluated for other genes.

## Description of Supplemental Data

Supplemental Data include notes on data collection from Stanford internal resources (CF Center and MPL) and four tables.

## Acknowledgements

This work was supported by the ClinGen grant number U01 HG007436-01. AR would like to acknowledge additional support from The Dean’s Postdoctoral Fellowship at the Stanford School of Medicine (2014-2015), and CEHG Postdoctoral Fellowship (2014-2015).

## Web Resources

Complete list of 1,899 protein-coding *CFTR* variants with clinical significance, allele frequency and corresponding source dataset name, as well as 35 annotation features for 1,210 missense variants together with RF predictions, are available on the GitHub: https://github.com/rychkova/CFTR-MetaPred
.

## Supplemental Data

### On the data collection for the Stanford CF center

The Stanford CF Center provides state-of-the-art care for a patient population that comes from the San Francisco Bay area and beyond. The center currently cares for approximately 450 patients, including post-transplant CF patients. Our current standing protocol for clinical care includes detailed phenotypic characterization as well as complete *CFTR* mutation analysis in all the patients under follow up. Information of *CFTR* mutations identified and results of sweat chloride results is kept on a secure database managed by the Center.

### On the data collection for Stanford MPL

The Stanford Molecular Pathology Laboratory provides clinical diagnostic testing for CF. The laboratory currently offers carrier screening (basic and expanded), diagnostic testing, and molecular testing associated with CF newborn screening for the State of California. This testing includes a 40 mutation screening panel for all newborns in the State of California who have a high initial IRT enzyme test result. A subset of these, namely those for whom only one mutation was identified through panel testing, receives further screening by direct DNA sequencing.

### Supplemental Tables

**Table S1.** Known CFTR motifs responsible for protein-protein interaction.

| **PPI motif** | **motif description** | **residues in the motif** | **reference** |
| --- | --- | --- | --- |
| SNAREs_binding_motif | Mediates membrane fusion and vesicle trafficking by assembling into complexes that link vesicle-associated SNAREs (v-SNAREs) with SNAREs on target membranes (t-SNAREs). | 1-79 | ref<sup>37</sup> |
| AMPK_binding_motif | A molecular switch that links ion transport to cellular metabolism. AMPK is activated when the AMP/ATP ratio increases. | 1420-1457 | ref<sup>38</sup> |
| PP2A_binding_motif | A heterotrimeric protein phosphatase that interacts with and dephosphorylates CFTR. | 1451-1476 | ref<sup>37</sup> |
| PDZ_binding_motif | Mediate protein-protein interactions by binding to short peptide sequences that are most often in the C termini of the target proteins. | 1477-1480 | ref<sup>39</sup> |
| endocytic_motif | Regulates endocytosis by which extracellular material and membrane-resident proteins are taken up by cells. | 1424-1427, 1430-1431 | ref<sup>37,38,40</sup> |
| possible_endocytic_motif | *Possible internalization signal suggested by R31C & R31L mutation experiments<sup>40</sup>.<br>**Possible internalization motif based on the structural similarity to conventional internalization motifs in NBD2. | 28-31*, 627-630**, 633-634** | ref<sup>40</sup> |

**Table S2.** Importance for all 35 features on the scale 0-100.

| **feature name** | **feature class** | **importance** |
| --- | --- | --- |
| AF | derived from datasets | 100.0 |
| Density | derived from datasets | 70.2 |
| MutPred | sequence & structure-based | 54.3 |
| POSE | sequence & structure-based | 31.6 |
| SIFT | sequence-based | 18.1 |
| topology | derived from protein structure | 17.7 |
| PPI_score | derived from protein structure | 14.7 |
| volume_change | physicochemical property | 13.6 |
| CADD_RawScore | sequence & structure-based | 13.3 |
| Eris_BenTal | stability predictor | 12.8 |
| FoldX_BenTal | stability predictor | 12.4 |
| HB_resi_change | derived from protein structure | 12.1 |
| nucleotide_binding | derived from protein structure | 11.4 |
| domain | derived from protein structure | 11.1 |
| SAA_BenTal | derived from protein structure | 10.0 |
| HB_total_change | derived from protein structure | 10.0 |
| PolyPhen2 | sequence & structure-based | 9.9 |
| Eris_Dokholyan | stability predictor | 9.3 |
| PoPMuSiC_Dokholyan | stability predictor | 9.2 |
| CADD_PHRED | sequence & structure-based | 7.6 |
| polarity_change | physicochemical property | 7.6 |
| modifications | derived from protein structure | 7.0 |
| PoPMuSiC_BenTal | stability predictor | 6.7 |
| PROVEAN | sequence-based | 6.6 |
| ch_contact_resi | derived from protein structure | 5.2 |
| SAA_Dokholyan | derived from protein structure | 5.1 |
| PANTHER | sequence-based | 4.4 |
| SAA_total_change | derived from protein structure | 4.3 |
| partition_energy_change | physicochemical property | 3.6 |
| hydrophobicity_change | physicochemical property | 3.0 |
| mem_contact_resi | derived from protein structure | 2.6 |
| SAA_resi_change | derived from protein structure | 2.1 |
| charge_change | physicochemical property | 1.5 |
| pro_signal | physicochemical property | 0.5 |
| FoldX_Dokholyan | stability predictor | 0.0 |

**Table S3.** Experimental *in vitro* mean Cl^-^ conductance measures for various *CFTR* missense variants.

| **aachange** | **clin_sign** | **mean_CI_cond** | **RF.prob** | **RF.pred** |
| --- | --- | --- | --- | --- |
| G576A | benign | 147.0 | 0.61 | pathogenic |
| I148T | benign | 91.1 | 0.17 | benign |
| L997F | benign | 22.4 | 0.33 | benign |
| R1162L | benign | 130.2 | 0.31 | benign |
| R31C | benign | 105.3 | 0.17 | benign |
| R668C | benign | 57.6 | 0.09 | benign |
| R75Q | benign | 65.4 | 0.16 | benign |
| S1235R | benign | 78.7 | 0.23 | benign |
| A455E | pathogenic | 6.8 | 0.92 | pathogenic |
| A559T | pathogenic | 0.0 | 1.00 | pathogenic |
| D110H | pathogenic | 9.1 | 0.92 | pathogenic |
| D614G | pathogenic | 18.0 | 0.92 | pathogenic |
| E92K | pathogenic | 0.2 | 0.90 | pathogenic |
| G1244E | pathogenic | 1.0 | 0.96 | pathogenic |
| G178R | pathogenic | 3.4 | 0.97 | pathogenic |
| G551D | pathogenic | 1.3 | 0.96 | pathogenic |
| G85E | pathogenic | 2.4 | 0.97 | pathogenic |
| G970R | pathogenic | 2.8 | 0.95 | pathogenic |
| I1234V | pathogenic | 39.9 | 0.84 | pathogenic |
| I336K | pathogenic | 0.9 | 0.91 | pathogenic |
| L1065P | pathogenic | 0.0 | 0.96 | pathogenic |
| L1077P | pathogenic | 0.0 | 0.94 | pathogenic |
| L206W | pathogenic | 5.7 | 0.97 | pathogenic |
| L227R | pathogenic | 0.0 | 0.69 | pathogenic |
| L467P | pathogenic | 0.0 | 0.92 | pathogenic |
| L927P | pathogenic | 0.1 | 0.93 | pathogenic |
| M1101K | pathogenic | 0.0 | 0.81 | pathogenic |
| M1V | pathogenic | 0.7 | 0.93 | pathogenic |
| N1303K | pathogenic | 0.5 | 0.90 | pathogenic |
| P205S | pathogenic | 2.5 | 0.93 | pathogenic |
| P67L | pathogenic | 6.0 | 0.87 | pathogenic |
| R1066C | pathogenic | 0.0 | 0.71 | pathogenic |
| R1066H | pathogenic | 1.3 | 0.82 | pathogenic |
| R117C | pathogenic | 3.4 | 0.84 | pathogenic |
| R334W | pathogenic | 1.3 | 0.92 | pathogenic |
| R347H | pathogenic | 5.2 | 0.97 | pathogenic |
| R347P | pathogenic | 0.0 | 0.94 | pathogenic |
| R352Q | pathogenic | 3.1 | 0.89 | pathogenic |
| R560K | pathogenic | 0.0 | 0.99 | pathogenic |
| R560T | pathogenic | 0.1 | 0.98 | pathogenic |
| S1251N | pathogenic | 5.2 | 0.93 | pathogenic |
| S341P | pathogenic | 0.0 | 0.88 | pathogenic |
| S492F | pathogenic | 0.0 | 0.93 | pathogenic |
| S549N | pathogenic | 1.6 | 0.90 | pathogenic |
| S549R | pathogenic | 0.1 | 0.96 | pathogenic |
| S945L | pathogenic | 6.0 | 0.96 | pathogenic |
| T338I | pathogenic | 1.0 | 0.66 | pathogenic |
| V520F | pathogenic | 0.2 | 0.97 | pathogenic |
| Y569D | pathogenic | 0.0 | 0.95 | pathogenic |
| D1152H | VUS | 77.8 | 0.43 | benign |
| D1270N | VUS | 53.2 | 0.88 | pathogenic |
| D579G | VUS | 13.9 | 0.88 | pathogenic |
| F1052V | VUS | 87.0 | 0.84 | pathogenic |
| G1069R | VUS | 122.8 | 0.80 | pathogenic |
| I1027T | VUS | 111.3 | 0.69 | pathogenic |
| R1070W | VUS | 8.9 | 0.81 | pathogenic |
| R117H | VUS | 35.0 | 0.67 | pathogenic |
| R74W | VUS | 44.0 | 0.47 | benign |
| S977F | VUS | 6.7 | 0.77 | pathogenic |
| V754M | VUS | 140.0 | 0.57 | pathogenic |

Clin\_sign – clinical significance of variants, RF.prob and RF.pred – probability of pathogenicity and predicted pathogenicity class by RF model. Mean CI<sup>-</sup> conductance values and clinical significance are from ref<sup>30</sup>.

**Table S4.** Mean sweat Cl^-^ concentration and RF probabilities for missense variants in CF patients of The Stanford CF Center heterozygous for F508del mutation.

| **aachange** | **clin_sign** | **mean_sweat_Cl** | **RF.prob** | **RF.pred** |
| --- | --- | --- | --- | --- |
| S1235R | benign | 19.0 | 0.23 | benign |
| D836Y | pathogenic | 15.0 | 0.70 | pathogenic |
| E217G | pathogenic | 24.0 | 0.65 | pathogenic |
| G194V | pathogenic | 112.0 | 0.79 | pathogenic |
| G27R | pathogenic | 96.5 | 0.73 | pathogenic |
| G551D | pathogenic | 98.1 | 0.96 | pathogenic |
| G85E | pathogenic | 112.0 | 0.97 | pathogenic |
| L1065P | pathogenic | 130.0 | 0.96 | pathogenic |
| N1088D | pathogenic | 69.0 | 0.88 | pathogenic |
| N1303K | pathogenic | 112.0 | 0.90 | pathogenic |
| R117C | pathogenic | 108.0 | 0.84 | pathogenic |
| R334W | pathogenic | 104.0 | 0.92 | pathogenic |
| R347H | pathogenic | 72.5 | 0.97 | pathogenic |
| R352W | pathogenic | 34.5 | 0.90 | pathogenic |
| R560T | pathogenic | 160.0 | 0.98 | pathogenic |
| S492F | pathogenic | 76.0 | 0.93 | pathogenic |
| S945L | pathogenic | 88.0 | 0.96 | pathogenic |
| V456A | pathogenic | 74.0 | 0.82 | pathogenic |
| V520F | pathogenic | 120.0 | 0.97 | pathogenic |
| V520I | pathogenic | 36.0 | 0.92 | pathogenic |
| V562I | pathogenic | 14.0 | 0.90 | pathogenic |
| D1152H | VUS | 38.0 | 0.43 | benign |
| F1052V | VUS | 41.0 | 0.84 | pathogenic |
| I1027T | VUS | 70.3 | 0.69 | pathogenic |
| R1070W | VUS | 23.0 | 0.81 | pathogenic |
| R117H | VUS | 66.6 | 0.67 | pathogenic |

Clinical significance of variants is based on The Stanford CF Center data.

